# Flower Power: Floral reversion as a viable alternative to nodal micropropagation in *Cannabis sativa*

**DOI:** 10.1101/2020.10.30.360982

**Authors:** A.S. Monthony, S. Bagheri, Y. Zheng, A.M.P. Jones

## Abstract

The legalization of *Cannabis sativa* L. for recreational and medical purposes has been gaining global momentum, leading to a rise in interest in *Cannabis* tissue culture as growers look for large-scale solutions to germplasm storage and clean plant propagation. Mother plants used in commercial propagation are susceptible to insect pests and disease and require considerable space. While micropropagation can produce disease free starting material in less space, current published *in vitro* micropropagation methods are not robust and few report high multiplication rates. Further, these micropropagation methods rely on photoperiod-sensitive plants which can be maintained in a perpetual vegetative state. Current methods are not adaptable to long-term tissue culture of day-neutral cultivars, which cannot be maintained in perpetual vegetative growth. In this study, we chose to develop a micropropagation system which uses *C. sativa* inflorescences as starting materials. This study used two cannabis cultivars, two plant growth regulators (PGR; 6-benzylaminopurine and meta-topolin) at different concentrations, and two different numbers of florets. Here we show that floral reversion occurs from meristematic tissue in *C. sativa* florets and that it can be used to enhance multiplication rates compared to existing *in vitro* methods. Floret number was shown to have a significant impact on percent reversion, with pairs of florets reverting more frequently and producing healthier explants than single florets, while cultivar and PGR had no significant effect on percent reversion. Compared with our previously published nodal culture studies, the current floral reversion method produced up to eight times more explants per tissue culture cycle. Floral reversion provides a foundation for effective inflorescence-based micropropagation systems in *C. sativa*.

## Introduction

Tissue culture of *Cannabis sativa* L. has become a topic of increasing interest as Canada and many US jurisdictions have legalized recreational and/or medicinal uses of *Cannabis*, mirroring a broader global trend. Micropropagation of *Cannabis* offers several advantages over traditional vegetative propagation methods used by licensed producers operating in greenhouses and indoor growth facilities. *In vitro* storage of germplasm is highly space efficient compared to maintaining mother plants for stem cuttings, which can occupy up to 15% of available floor space in a commercial growth facility. In addition, propagation and maintenance of photoperiod-insensitive cultivars of *Cannabis* under normal greenhouse conditions is especially challenging as they cannot be maintained in a perpetual vegetative state required to maintain mother plants (Piunno et al. 2019). *In vitro* growth systems also offer a source of axenic, disease-free tissues and are a pre-requisite to the implementation of plant biotechnologies. *Cannabis* micropropagation protocols have largely been developed using meristematic tissues from axillary or apical nodes (Richez-Dumanois et al. 1986; Ślusarkiewicz-Jarzina et al. 2005; Plawuszewski et al. 2006; Lata et al. 2009b, a, 2016; Smýkalová et al. 2019; Wróbel et al. 2020; Page et al. 2020). While some publications report promising accounts of multiple shoot cultures (MSCs) from nodal tissues (Lata et al. 2009b, a, 2016), low levels of shoot proliferation rates and multiplication rates are more common (Richez-Dumanois et al. 1986; Ślusarkiewicz-Jarzina et al. 2005; Plawuszewski et al. 2006; Smýkalová et al. 2019; Wróbel et al. 2020; Page et al. 2020). The limited success of MSCs have been attributed in part to strong apical dominance in the shoots which reduces branching and multiple stem formation (Smýkalová et al. 2019; Wróbel et al. 2020). Recent work has highlighted the concerns surrounding the replicability and the variability between published works in the area of *C. sativa* tissue culture (Monthony et al. 2020b; Page et al. 2020).

One alternative to nodal culture is the use of *Cannabis* inflorescences as starting material for multiplication. Floral reversion is a process in which floral tissues revert to a vegetative state and has been demonstrated *in vitro* across species from many taxa (Eapen and George 1997; Phulwaria and Shekhawat 2013; Punyarani et al. 2013; Shareefa et al. 2019). In the development of cell suspension cultures, Raharjo *et al*. (2006) found that *Cannabis* flowers were more responsive to callus formation than leaves; however, the regenerative potential of flowers was not investigated in this study. A recent publication by Piunno *et al*. (2019) provided the first known report of *in vitro* shoots recovered from floral explants of *C. sativa*. In this study Piunno *et al*. (2019) achieved low rates of vegetative shoot production from floret clusters sourced from greenhouse-grown inflorescences in two high-tetrahydrocannabinol (THC) cultivars of *C. sativa*. These results were the first to show putative floral reversion of *Cannabis* inflorescences. The cellular development which characterizes floral reversion have been differentially described depending on the species studied (Zayed et al. 2016). In some species floral reversion occurs from existing meristems (Sen et al. 2013) while from others reversion is a result of *de novo* regeneration (Poluboyarova et al. 2014; Zayed et al. 2016). The mode of floral reversion has implications in plant biotechnology where early transient gene expression studies have shown the potential to express transgenes in floral tissues (Deguchi et al. 2020). Despite observing reversion and normal subsequent growth of vegetative plants, Piunno *et al*. (2019) did not identify the developmental pattern of floral reversion in *Cannabis*.

Existing literature has demonstrated the successful use of floral reversion in commercially important crops and recalcitrant species. *In vitro* floral reversion has been used to overcome challenges faced by traditional multiplication protocols, which can be limited by low multiplication rates, high levels of contamination and a limited amount of suitable plant material for propagation (Gubišová et al. 2013; Zayed et al. 2016; Shareefa et al. 2019). In many cases, inflorescences have been more successful than other explant sources, making floral reversion an important tool to overcoming recalcitrance and for the rapid production of clonal, disease-free elite-cultivars (Zayed et al. 2016; Shareefa et al. 2019). While many plants do not flower *in vitro*, a recent study by Moher *et al*. (2020) found that flowering rates exceeded 75% under a 12-hour photoperiod in *in vitro* grown *Cannabis* explants. The authors also found that flowering occurred rapidly, with a majority of explants flowering in under 20 days (Moher et al. 2020). Considering the strong flowering response observed *in vitro* and the challenges of replicability, multiplication rate and apical dominance experience using *in vitro* nodal cuttings, we hypothesized that floral reversion using inflorescences from *in vitro* plants can provide an alternative approach to vegetative micropropagation of *C. sativa*.

The objective of this study was to: I) Optimize floral reversion for micropropagation by comparing the effect of floret number and cytokinins on the efficacy of the process and; II) Identify the source of tissues responsible for reversion through histological sampling. In this study we show that pairs of florets increase the probability of reversion, and result in larger explants and that reversion occurs from existing meristem regions in the inflorescences. Based on high florets density of *C. sativa* inflorescences, we propose that floral reversion can improve the number of plants produced per tissue culture cycle when compared with nodal shoot proliferation methods.

## Materials & Methods

### Plant Material

*In vitro* maintained plants of two high-THC cultivars of *C. sativa*, U82 and U91 (Hexo Corp.; Monthony et al., 2020) were used as sources of *in vitro* florets. These were maintained on DKW basal medium as reported by Page et al. (2020). To induce flowering, *in vitro* explants maintained in GA-7 vessels (Magenta, Illinois) were transferred to a 12/12 (dark/light) photoperiod. Plants were grown under Photoblasters (WeVitro Inc., ON) ~50 μmol s^−1^ m^−2^ as described by Moher *et al*. (2020) for ~ 2 months at 25 °C prior to floret dissection. Single (Fig. 1A) and pairs (Fig. 1D) of florets (hereafter referred to as ‘singles’ and ‘pairs’) were dissected using scalpel and forceps under axenic conditions using a digital dissecting microscope (Koolertron, China) in a laminar flow hood (Design Filtration Microzone, ON).

**Fig. 1.**
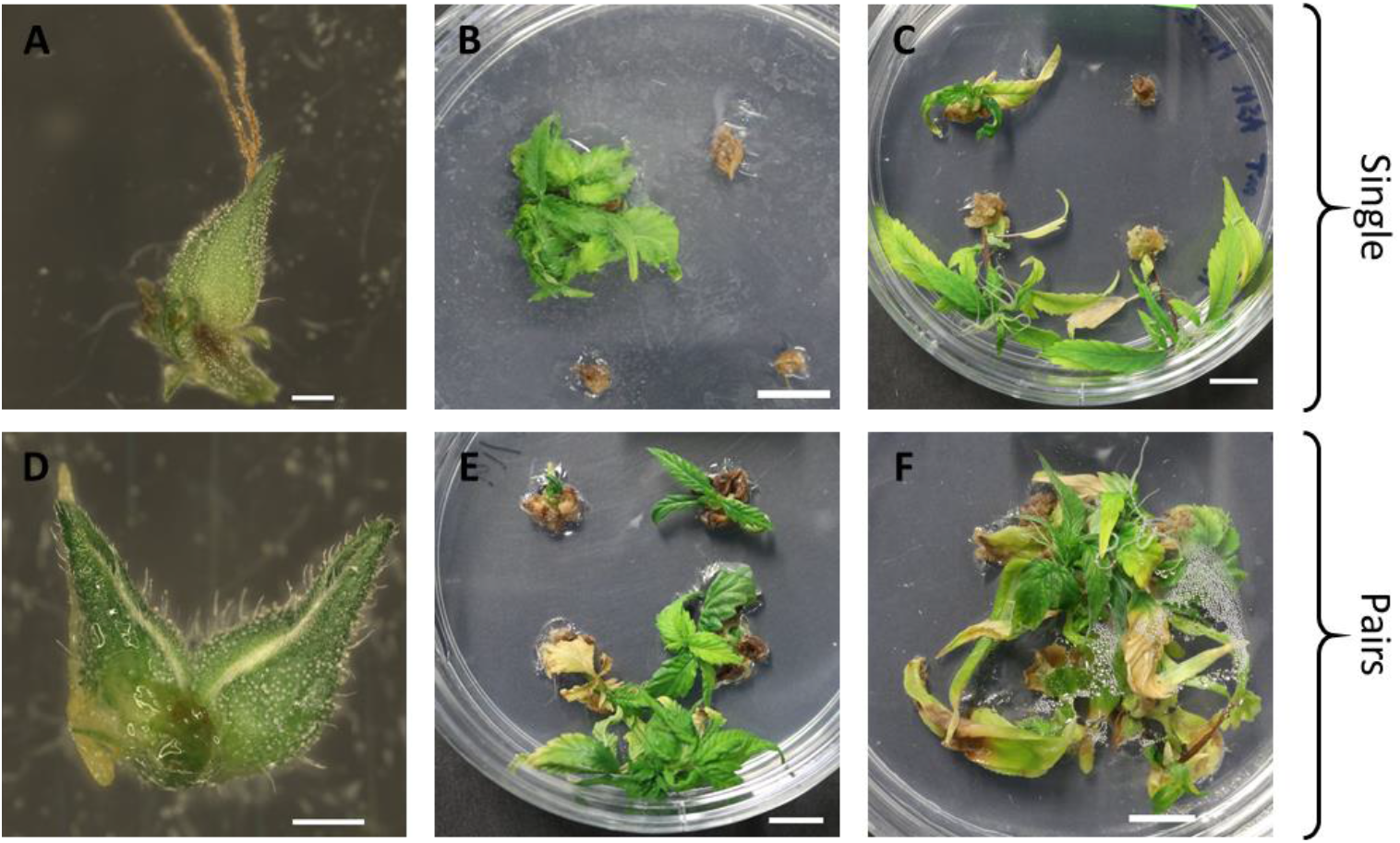
Single (A) and pair (D) of freshly dissected (day 0) florets from *Cannabis* cv. U91. Scale bar 1 mm. B) 8 week single florets from U91 cultured on DKW media with 1 μM BAP. C) 6 week single florets from U91 cultured on DKW media with 1 μM mT. E) 8 week pairs of florets from U91 cultured on DKW media with 1 μM BAP. F-6 week pairs of florets from U91 cultured on DKW media with 1 μM mT. Scale bar for B, C, E and F is 1 cm.

### Media preparation

Media consisted of DKW Salts with Vitamins (D2470, PhytoTech Labs, KS), 3% sucrose (w/v), 0.6% agar (w/v) (Fisher Scientific) dissolved in distilled water and pH was adjusted to 5.7 using 1 N NaOH. Media were amended with plant growth regulators (PGRs) from 1 mM stock solutions stored at 4 °C for a final concentration of 0, 0.01, 0.1, 1.0 and 10 μM of either 6-benzylaminopurine (BAP) or meta-topolin (mT). BAP was added prior to pH adjustment and autoclaving. Due to its heat labile nature, mT was added via filter sterilization to media after it was autoclaved and cooled to ~60 °C. The media were autoclaved for 20 minutes at 121 °C and 18 PSI. Approximately 25 mL of the autoclaved media were dispensed into sterile 100 x 20 mm Petri dishes (VWR International, ON) in a laminar flow hood (Design Filtration Microzone).

### Experimental design

To assess the efficacy of BAP on reversion in single and pairs of florets in both *C. sativa* - cultivars (U82 and U91) a 2×2×5 cross-classified factorial experiment with a completely randomized design with three factors was used. The main effects were A) cultivar (U82 and U91); B) concentration of BAP in μM (0.0, 0.01, 0.1, 1.0 and 10); and C) floret number (single florets vs. pairs of florets). Each treatment consisted of 4 experimental units (Petri dishes; *n=4*), each with 4 floral explants (sampling units). The explants (floret pairs or singles) were maintained on the prepared semi-solid media containing the appropriate concentration of BAP in a controlled atmosphere walk-in growth chamber, under LED lighting (Fig. S1) using a 16-hr photoperiod at 25 °C with a light intensity of ~ 50 μmol s^−1^ m^−2^ measured using a LI-180 Spectrometer (LI-COR, NE), until reverted vegetative explants began to show signs of crowding in the Petri dishes (8 weeks).

**Fig. S 1.**
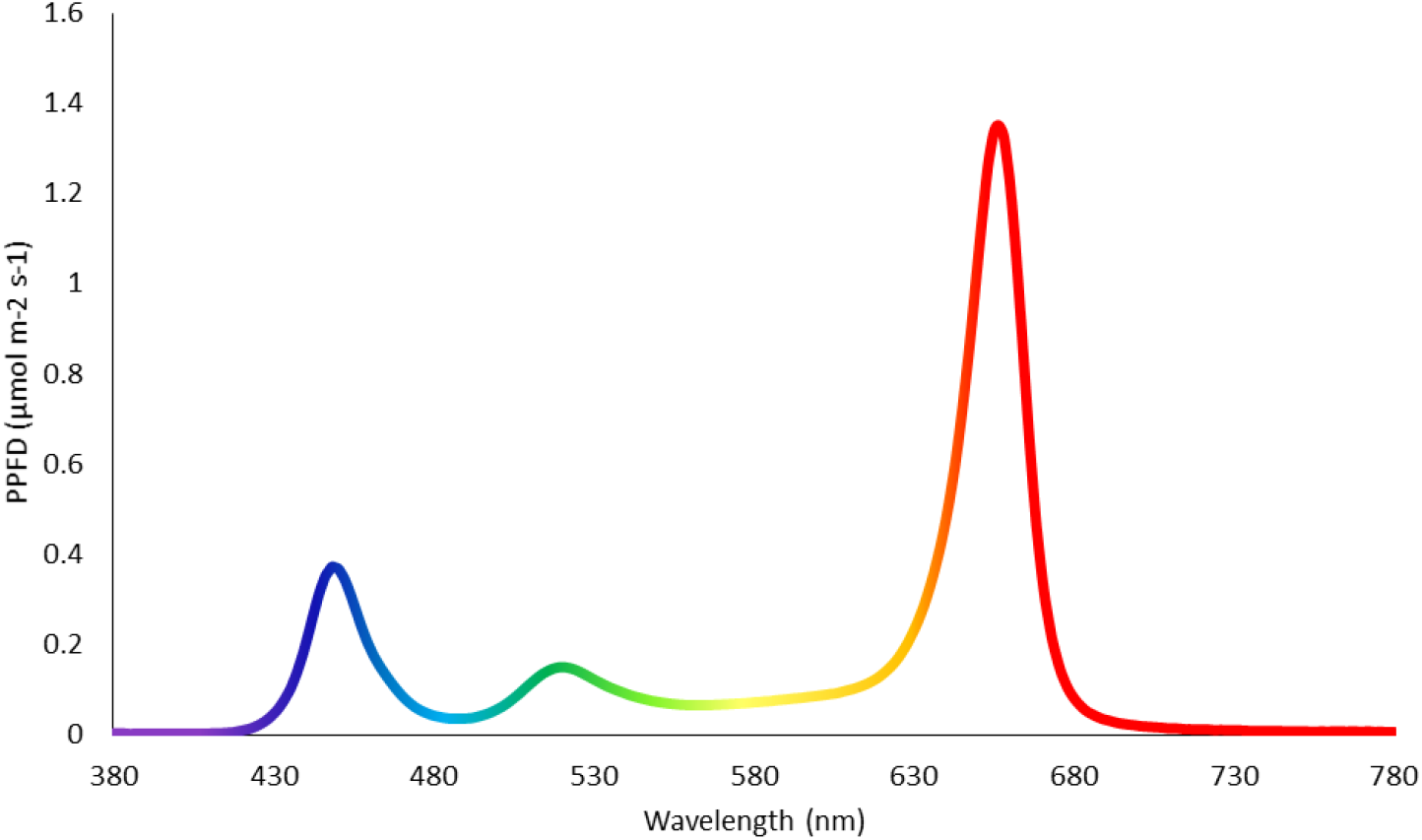
A sample spectrum of the LED lighting used to maintain reverting explants, obtained using a LI-COR LI-180 Spectrometer. PPFD: Photosynthetic Photon Flux Density.

The second experiment assed the effect of mT on reversion in single and pairs of florets in *C. sativa* cv. U91. The experiment was a 2×5 cross-classified factorial experiment with a completely randomized design with two factors. The main effects were concentration of mT in μMolar (0.0, 0.01, 0.1, 1.0 and 10), and floret number. The explants were maintained on the prepared semi-solid media containing mT in a controlled atmosphere walk-in growth chamber as previously described, until reverted vegetative explants began to show signs of crowding in the Petri dishes (6 weeks).

The average number of healthy florets which could be isolated was determined by counting the number of healthy florets dissected from 11 flowering explants. Explant fresh weight was determined at the end of the experiment by removing the initial florets and any callus formed on the reverted explant and then weighing each reverted explant on a balance (Quintix^®^ 2102-1S, Sartorius, Germany). Response rate was determined by counting the number of florets that reverted and dividing by the total number of florets on each Petri dish (a floret pair is considered as one explant). To obtain canopy area per explant, each Petri dish was photographed at the end of the experiment and the canopy area of each explant was determined by dividing the total canopy area in the photograph by the number of responsive explants. Canopy area measurement were performed using ImageJ software (v1.53a, National Institute of Mental Health, MD, USA). In short, images were cropped using the freehand selection tool to only include responding explants. The image background was removed (Process> Subtract background). Canopy area was selected by colour thresholding for green (Image>Adjust>Color Threshold) and measured (Analyze>Measure). Shoot proliferation rate was determined by dividing the number of vegetative explants subcultured from each Petri dish by the number of floral explants which underwent reversion in each Petri dish. A floral multiplication index, representing the average number of plants produced per cycle of micropropagation, was calculated using the average number of florets produced on a flowering explant, the average shoot proliferation rate and the average percentage reversion for each treatment using the following equation:

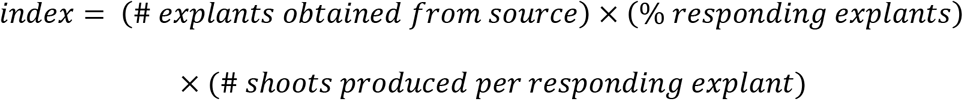

**Equation I-** Calculation of the floral multiplication index

### Rooting and *ex vitro* acclimatization

Following the reversion period, explants were transferred from Petri dishes to GA-7 vessels (Magenta) with approximately 50 mL of basal DKW medium consisting of DKW Salts with Vitamins (D2470, PhytoTech Labs, KS), 3% sucrose (w/v), 0.6% agar (w/v) (Fisher Scientific) in distilled water and pH adjusted to 5.7 using 1 N NaOH. Explants were maintained under a vegetative photoperiod (16 h light; 8 h dark) at 25 °C with a light intensity of ~ 50 μmol s^−1^ m^−2^. Explants developed roots within 1 month of subculture. Proportion of explants rooted were determined by counting the number of explants with at least one root > 5 mm in the culture vessel and dividing by the total number of explants in each culture vessel.

### Histological sampling & microscopy

Freshly dissected and 7-day old floral explants maintained on DKW basal media were fixed in 10% buffered formalin (ACP Chemicals Inc.). Samples were subsequently processed for paraffin embedding following the Standard Protocol for Formalin-Fixed Paraffin Embedded Tissue (FFPE) at the Animal Health Laboratory, Guelph, Ontario, Canada following Winegard *et al*. (2014). Sectioning was performed at 4 μm and staining was done with methylene blue. Slides were observed under bright field using an Axio Zoom.V16 microscope (Carl Zeiss Microscopy GmbH, Germany), and images were acquired and processed using Zen 2.3 Blue Edition (Carl Zeiss Microscopy GmbH).

### Statistical analysis

To assess the efficacy of BAP at inducing reversion in single and pairs of florets in both *C. sativa* cultivars (U82 and U91) a 2×2×5 factorial experiment with a completely randomized design was used with the following statistical model:

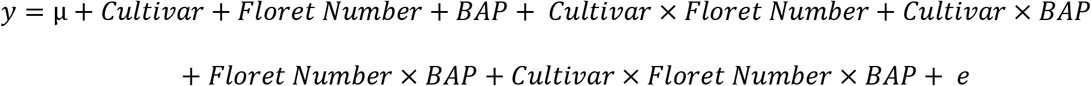

Where *y* is the measured response variables (response rate, canopy area, explant mass)

Where *μ* is the overall mean of the response variable

Where *Cultivar, Floret Number* and *BAP* are the fixed effects

Where *e* is the residual error

To assess the efficacy of mT at inducing reversion in single and pairs of florets *C. sativa* cv. U91 a 2×5 factorial experiment with a completely randomized design was used with the following statistical model:

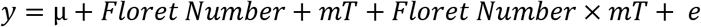

Where *y* is the measured response variables (response rate, canopy area, explant mass)

Where *μ* is the overall mean of the response variable

Where *Floret Number* and *mT* are the fixed effects

Where *e* is the residual error

All data were analyzed using SAS University Edition (SAS Studio 3.8; SAS Institute Inc.) Residual analyses were carried out for all response variables to check for normality. Canopy area and explant mass were normally distributed datasets and did not require subsequent adjustments prior to analysis. Due to a very narrow distribution of the data for multiplication rate, all data were lognormalized prior to analysis, and subsequently backtransformed after analysis. Prior to analysis, the percentage reversion data were transformed using an arcsine square root transformation, this was chosen over a logit transformation as the datasets for response were highly clustered at 0 and 1, and the arcsine square root model shows less dramatic variance at the end of the distribution. Rooting frequency (%) was analyzed using a binomial distribution (DIST= BINOMIAL, METHOD=QUAD) with a logit link to account for its non-Gaussian nature. An analysis of variance (ANOVA) was performed for all response variables using PROC GLIMMIX and means comparisons were obtained using the LSMEANS statement (α=0.05). Multiple comparisons were accounted for by a post-hoc Tukey-Kramer Test. Visual presentation of the SAS data and calculation of the % response means was done using Microsoft Excel^®^ (Microsoft Corp., WA, USA).

## Results

### Floral reversion

Floret number significantly affected the percent reversion of explants to the vegetative state in both the BAP and mT experiments (Table 1 and Table 2). In both experiments, pairs were approximately three times more likely to revert than single florets. The treatment average for all BAP treated explants (0 μM to 10 μM) found that 55% of floret pairs reverted compared to only 18% of single floret (*p* < 0.0001). The BAP treatment with the highest percentage reversion was 1 μM BAP using pairs of florets which achieved an average of 69% reversion (Fig. 2A). A similar trend was observed in mT treated of floral explants of cv. U91, where pairs of florets showed approximately 2.5 times higher rate of reversion (70% vs 28%; *p* < 0.0001) between the singles and pairs of floret treatment averages. Treatment of pairs of florets at 1 and 10 μM mT achieved the highest percentage reversion with 81% of florets reverting (Fig. 2B). While the percent reversion was significantly affected by the floret number, the number of shoots produced per explant was not significantly affected by any of the fixed effects, with each treatment producing between 1.5 and < 2 vegetative shoots per responding explant. Each flowering *in vitro* plant produced an average of 24 ± 6 healthy florets (or 12 pairs) for use in the experiments.

**Table 1.** Results of the F-test from the ANOVA for the percentage of florets which reverted (response variable). The 2×2×5 factorial designed tested the effects of cultivar, [BAP] and floret number and their interactions on the percentage of florets which underwent floral reversion.

| Fixed Effects | Numerator df | Denominator df | F Value | P-value |
| --- | --- | --- | --- | --- |
| Floret Number | 1 | 59 | 51.89 | <0.0001 |
| [BAP] | 4 | 59 | 2.05 | 0.0987 |
| Cultivar | 4 | 59 | 2.40 | 0.1265 |
| Floret Number x [BAP] | 1 | 59 | 2.06 | 0.0979 |
| Floret Number x Cultivar | 1 | 59 | 1.87 | 0.1766 |
| [BAP] x Cultivar | 4 | 59 | 1.81 | 0.1382 |
| Floret Number x [BAP] x Cultivar | 4 | 59 | 0.70 | 0.5968 |

**Table 2.** Results of the F-test from the ANOVA for the percentage of florets which reverted (response variable). The 2×5 factorial designed tested the effects of [mT] and floret number and their interactions on the percentage of florets which underwent floral reversion.

| Fixed Effects | Numerator df | Denominator df | F Value | P-value |
| --- | --- | --- | --- | --- |
| Floret Number | 1 | 30 | 27.50 | <0.0001 |
| [mT] | 4 | 30 | 2.57 | 0.0581 |
| Floret Number x [mT] | 4 | 30 | 0.95 | 0.4478 |

**Fig. 2.**
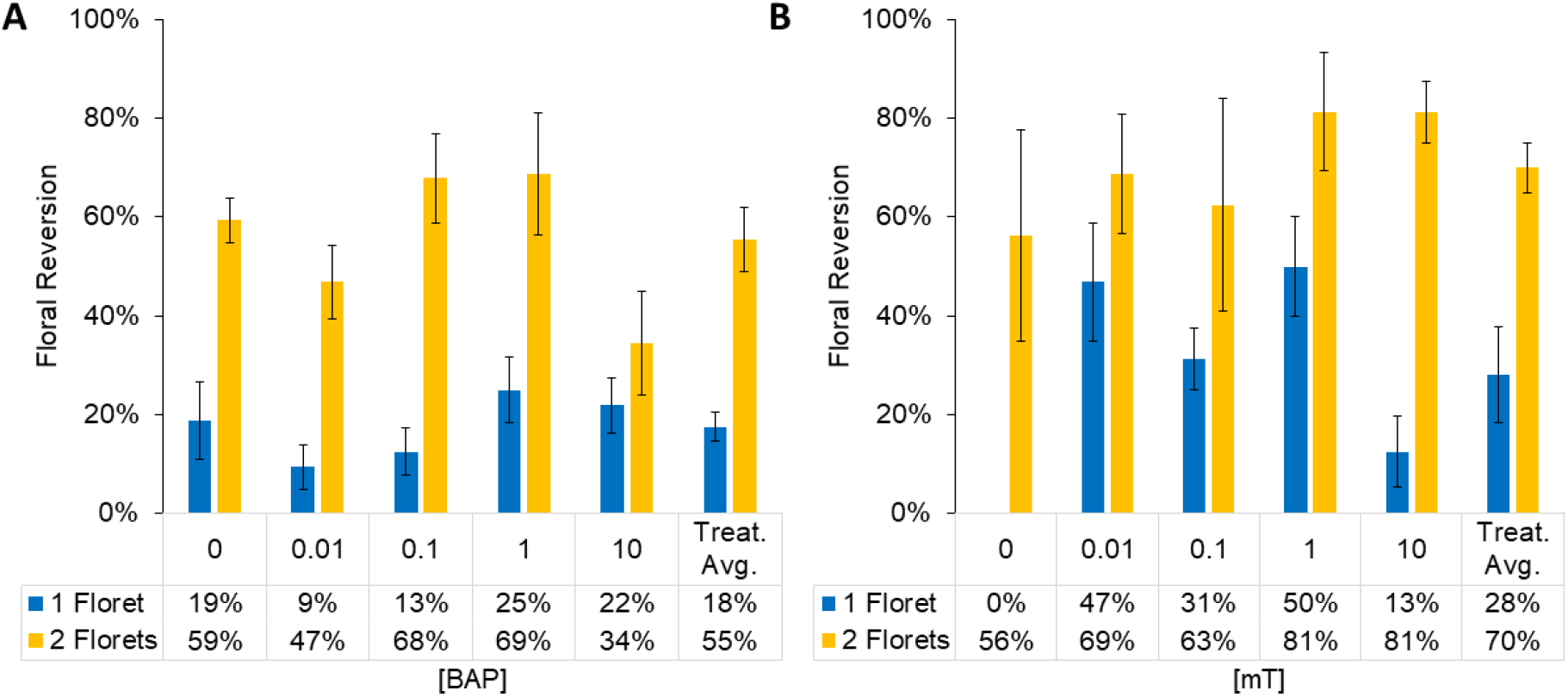
Percent reversion as a function of floret number and concentration of BAP (A) and mT (B). The interaction between Floret and [BAP] or [mT] were not significant as determined by an ANOVA (Table 1 and Table 2); however, 1 μM treatments showed the highest percentage of responding florets in both experiments. Percentage reversion reported in (A) is an average of both tested cultivars: U82, U91 where for each bar n=8. For mT only cv. U91 was tested, n=4. Treatment averages represent the means of 0-10 μM reversion percentages for single and for pairs of florets. Bars are mean ± standard error.

A multiplication index was calculated (Equation 1) using values from the treatment with the highest % reversion (1 μM mT) as a way of comparing the theoretical multiplication potential of this method with our previously published nodal shoot proliferation method (Page et al. 2020). The floral multiplication index represents the number of shoots that can be obtained (to transfer *ex vitro* or induce flowering to repeat the cycle) from a single flowering plant after one micropropagation cycle using floral reversion (Fig. 3). The values used to calculate the floral multiplication index (Equation 1) from the 1 μM mT treatment were: 51% reversion for singles and 81% for pairs, a shoot proliferation value of 1.5, and the average number of healthy florets produced per explant (24 singles/12 pairs). The floral multiplication index was calculated to be 18.2 plants per explant for single florets and 14.7 plants per explant for floret pairs at the 1 μM mT concentration.

**Fig. 3.**
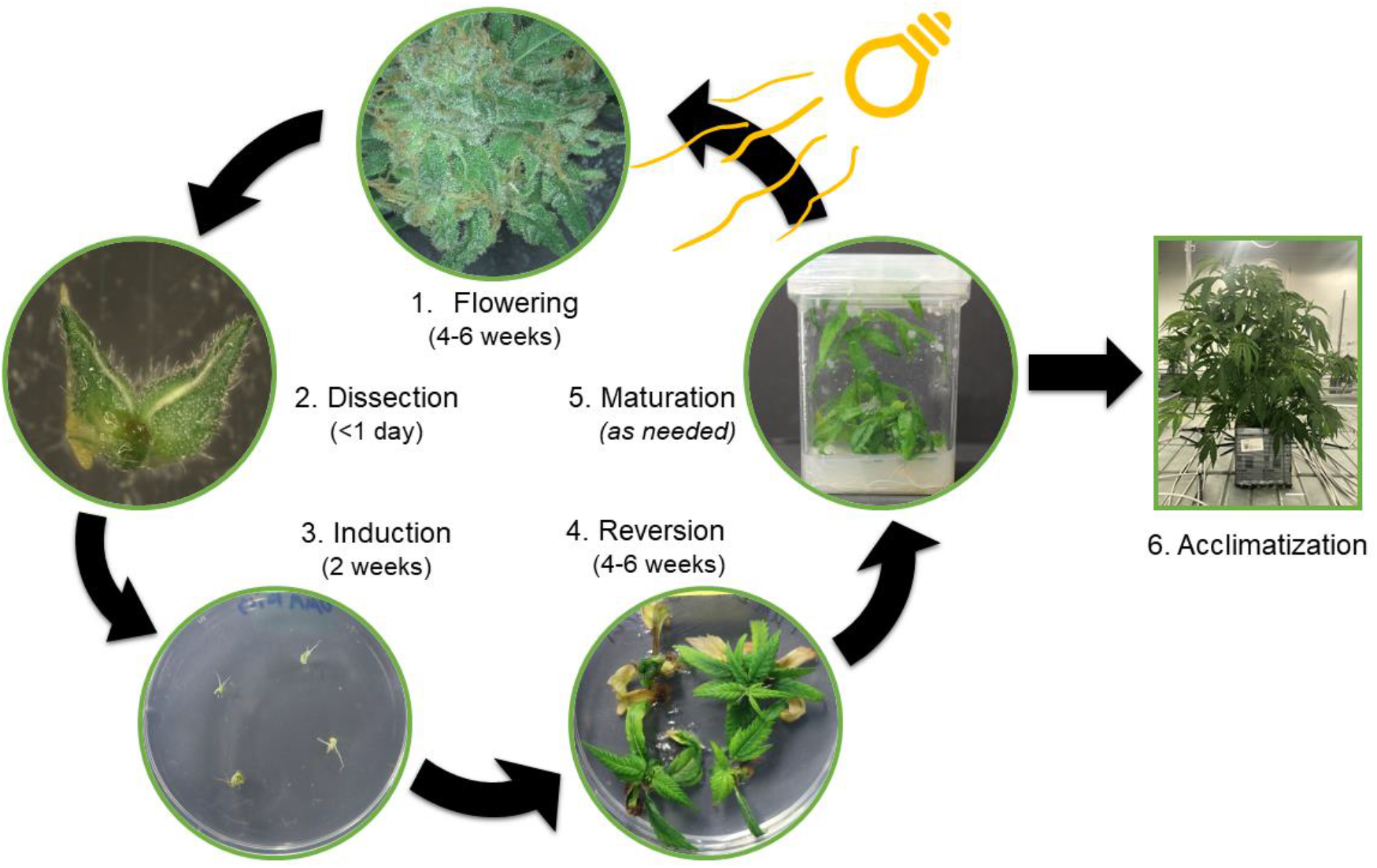
The proposed micropropagation cycle using floral reversion. 1. Flowering: Culture of mature vegetative explants under a 12-hour photoperiod to trigger *in vitro* flowering. 2. Dissection: Single or pairs of florets are dissected. 3. Induction: Florets are transferred to DKW-based media to begin floral reversion. 4. Reversion: Normal vegetative growth occurs, indicating reversion. 5. Maturation: Reverted explants cultured on DKW will root and can be used again for *in vitro* flowering or moved *ex vitro*. 6. Acclimatization: Reverted explants can be transferred to *ex vitro* conditions for hardening.

Floret number and the choice of PGR played a role in impacting the size of the reverted explants. In BAP treated explants, shoots that developed from pairs of florets produced larger canopy areas and greater fresh weights than shoots reverted from single florets (Fig. 4A). The average fresh weight of the BAP treated florets (0 μM to 10 μM) was 155 mg in singles compared to 232 mg in pairs (*p* = 0.0085; Fig. 4A). The difference in canopy area between singles and pairs in the BAP treated plants was similar to the difference in fresh weight with average areas of 1.12 cm^2^ for singles and 2.41 cm^2^ for doubles; (*p* = 0.0001; Fig. 4A). BAP treatment also significantly affected the explant mass (*p* = 0.0142) with 10 μM BAP resulting in an compared to the 0 μM control (255 mg vs. 138 mg) while 0.1 μM BAP resulted in a decrease in mass relative to the 0 μM control (125 mg vs. 138 mg). BAP alone, did not have a significant effect on canopy area (*p* = 0.0653). The interaction between the effects of floret number and BAP did not significantly affect the canopy area or explant mass (*p* > 0.05), however cultivar significantly affect the canopy area (*p* = 0.0069), with explants of U82 producing 1.6 times larger canopies compared to U91 (2.20 vs. 1.33 cm^2^ respectively). This difference in explant size did not, however, impact the *ex vitro* acclimatization, rooting and growth (Fig. 5).

**Fig. 4.**
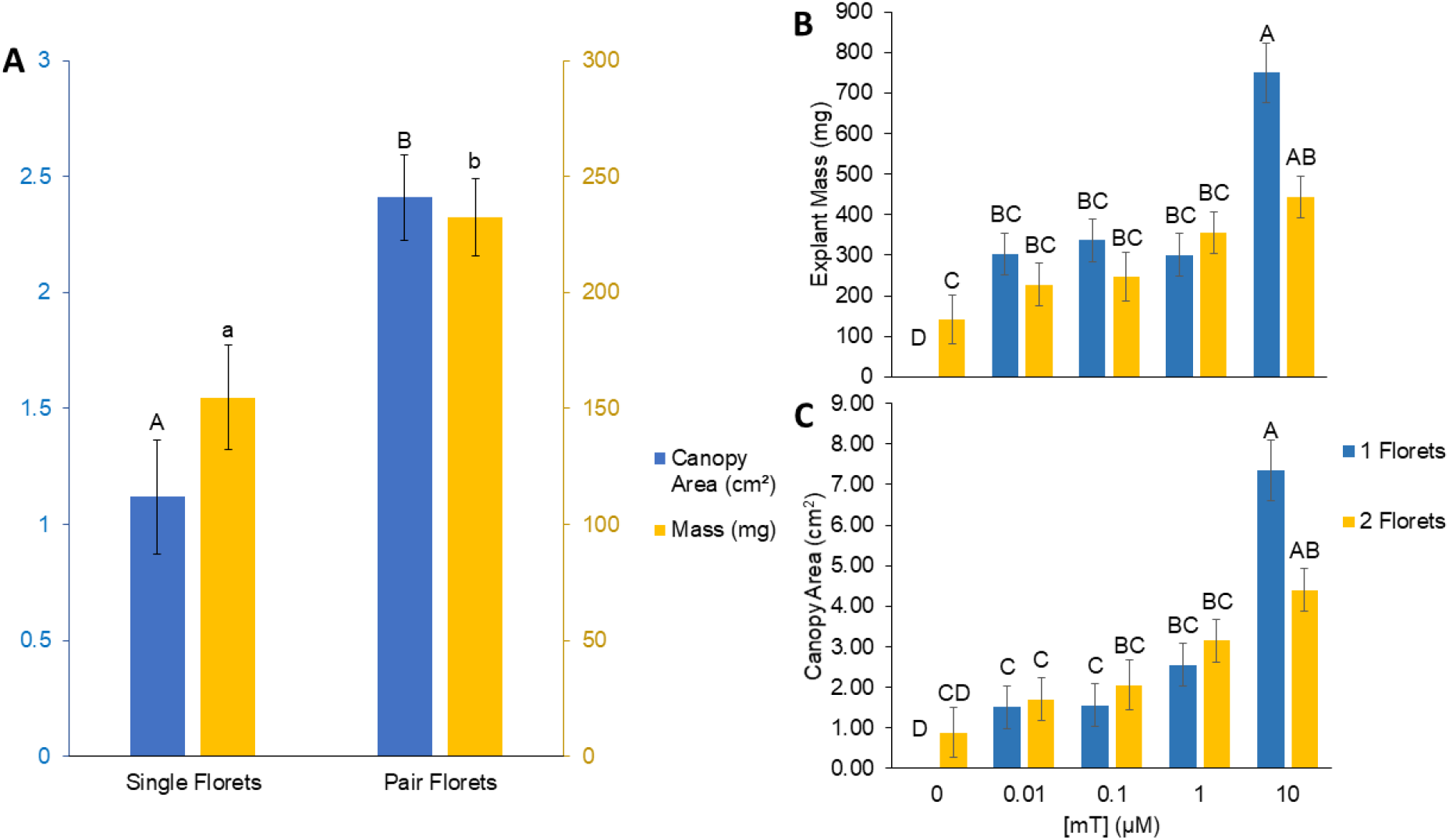
A) Fixed effect averages from BAP treated florets showing how both canopy and explant mass are significantly increased over 8 weeks in explants reverted from floret pairs. Bars bearing different letters are significant at *p* <= 0.01. Yellow bars correspond to mean explant mass across all BAP concentrations in milligrams and blue bars indicate mean canopy area across all BAP concentrations in cm^2^. Error bars are the standard error of the mean (n = 20). B) The significant interaction between floret number and mT concentration resulted in an increase in mass at higher mT concentrations after 6 weeks of culture. Error bars are the standard error of the mean (n = 4). C) The significant interaction between floret number and mT concentration also resulted in an increase in canopy area at higher mT concentrations after 6 weeks of culture. Error bars are the standard error of the mean. Bars bearing the same letter within either 1 Floret or 2 Florets are not significantly different at *p* < 0.05 as determined by a Tukey-Kramer multiple comparisons test (n=4).

**Fig. 5.**
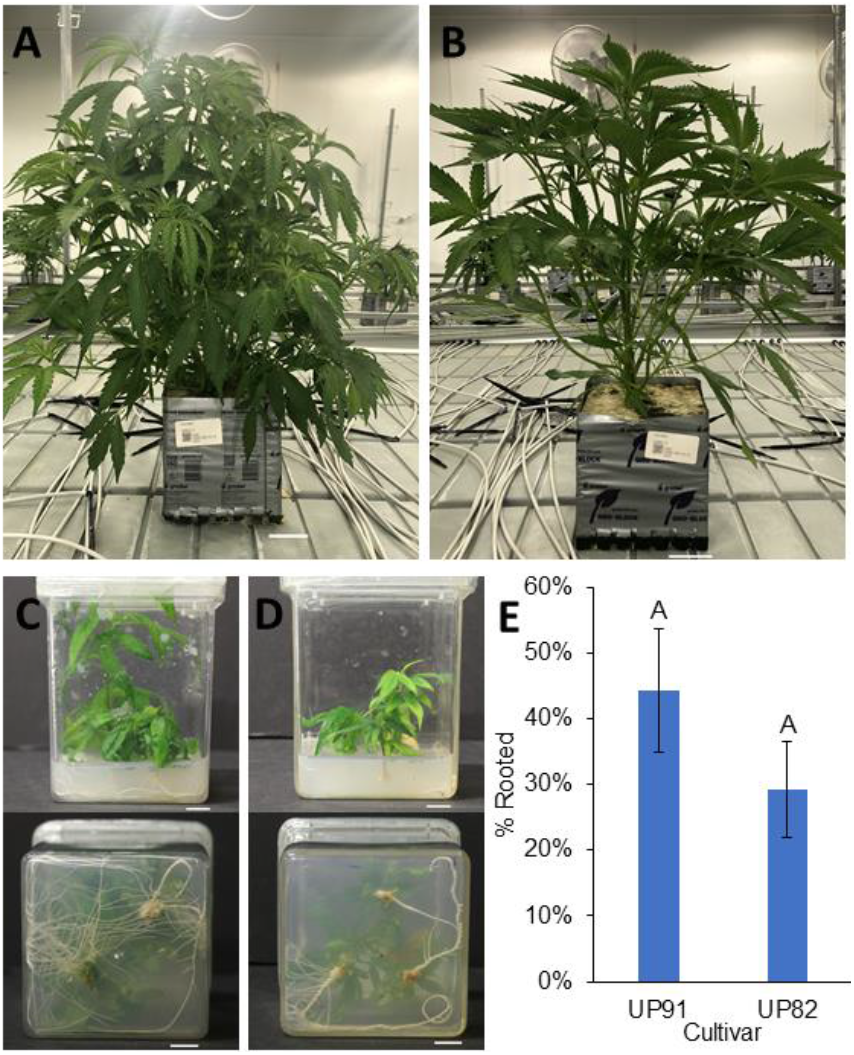
*In vitro* rooted plants were acclimatized and established *ex vitro* in commercial growth facilities in a soilless substrate. U82 (A) and U91 (B) are pictured after six weeks of *ex vitro* acclimatization. Images (A) and (B) were edited to remove confidential text information from the labels. Scale bar: 5 cm. A representative side and bottom view of U82 (C) and U91 (D) explants rooting *in vitro* on basal DKW media without auxin supplementation within 3 months of reversion. Scale bar: 1 cm. E) Percent rooting in the tested cultivars. Means with the same letter were not significantly different at *p* < 0.05 as determined by a Tukey-Kramer multiple comparisons test.

The mT and BAP trials were not run in tandem and as such, were not directly compared; however, qualitative observations found that mT leads to a more vigorous flush of initial growth in reverting explants compared to those grown on BAP. Earlier signs of crowding, stress and nutrient deficiency such as yellowing leaves, compared to BAP treatments (Fig. 1B, C, E and F and Fig. 6) resulted in mT treated explants being subcultured after 6 weeks (compared with 8 weeks for BAP treated explants) to avoid stunted growth or explant death. The interaction between floret number and mT significantly affected the explant mass (*p* = 0.0419; **Fig. 4B**) and the canopy area (*p* = 0.0239; Fig. 4C). The 10 μM mT treatment resulted in reverted explants with the largest masses: 750 mg from singles and 444 mg from pairs, compared to masses of 0 mg and 141 mg in the singles and pair controls treated with 0 μM mT (Fig. 4B). Despite a large explant mass and high % reversion at 10 μM mT, explant development showed more morphological abnormalities than the control: These explants were shorter and more compact with curled leaves and produced more callus than BAP treated explants (Fig. 6). Callogenesis was observed at 1 and 10 μM concentrations of BAP and mT; however, these calli were not organogenic and formed after the emergence of vegetative shoots (Fig. 6). Canopy area showed a similar trend as explant mass with 10 μM mT producing larger canopy areas per explant than the 0 μM mT control (Fig. 4C).

**Fig. 6.**
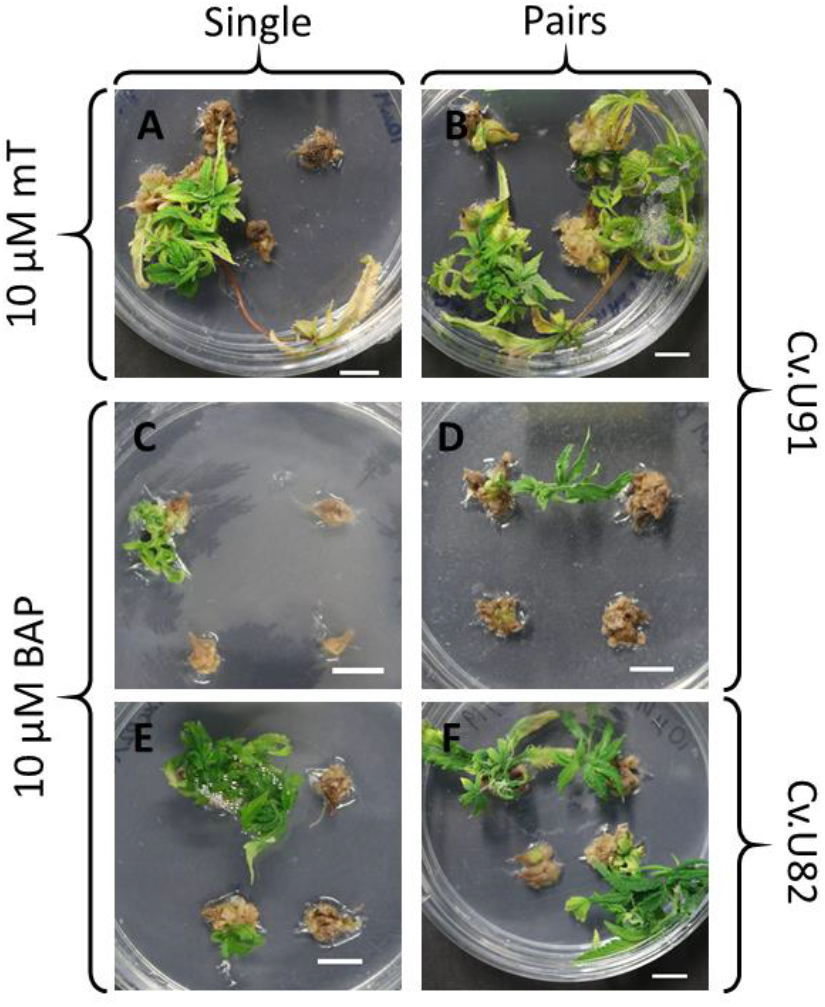
At 10 μM of BAP and of mT, morphology of the reverted vegetative explants was abnormal, with notable callus production after reversion had occurred. High concentrations of BAP impaired reversion, while high concentrations of mT caused vigorous but morphologically abnormal growth. A) 6 weeks single U91 at 10 μM mT B) 6 weeks pair U91 at 10 μM mT C) 8 weeks single U91 at 10 μM BAP D) 8 weeks pair U91 at 10 μM BAP E) 8 weeks single U82 at 10 μM BAP and F) 8 weeks pair U82 at 10 μM BAP. Scalebar 1 cm.

### Rooting and *ex vitro* acclimatization

Explants were transferred from the reversion media to basal DKW media without the inclusion of auxins once signs of nutrient deficiency and/or crowding of Petri dishes was observed (6 or 8 weeks). Both cultivars rooted with 44% of U91 and 29% of reverted U82 explants producing roots. Rooting percentage between the tested cultivars did not significantly differ (*p* = 0.1251). All explants transferred to *ex vitro* conditions were grown under T5 fluorescent lights at 150-200 μmol m^−2^ s ^−1^ with 50% relative humidity at 25 *°C* and showed normal morphological growth (Fig. 5 A & B).

### Histology

Histological sampling was performed to determine whether shoot proliferation was occurring *de novo* or from existing meristems. Samples were taken from freshly dissected florets (Fig. 7) and florets cultured on DKW basal media for 7 days (Fig. 8). Histological sampling of freshly dissected florets revealed the presence of meristem at their base (Fig. 7 C). Histological sampling of 7-day old floral undergoing reversion showed that this meristem region had undergone further development, revealing the presence of distinct vegetative meristems subtending the floral explant (Fig. 8). These vegetative meristems were flanked by floral meristems (Fig. 8 C & E) and were closely located to the excision area for the preparation of single florets used in this experiment (Fig. 7 C). The floral meristems which flank the vegetative meristems were further developed than the vegetative meristems that they flanked (Fig. 8 C & E).

**Fig. 7.**
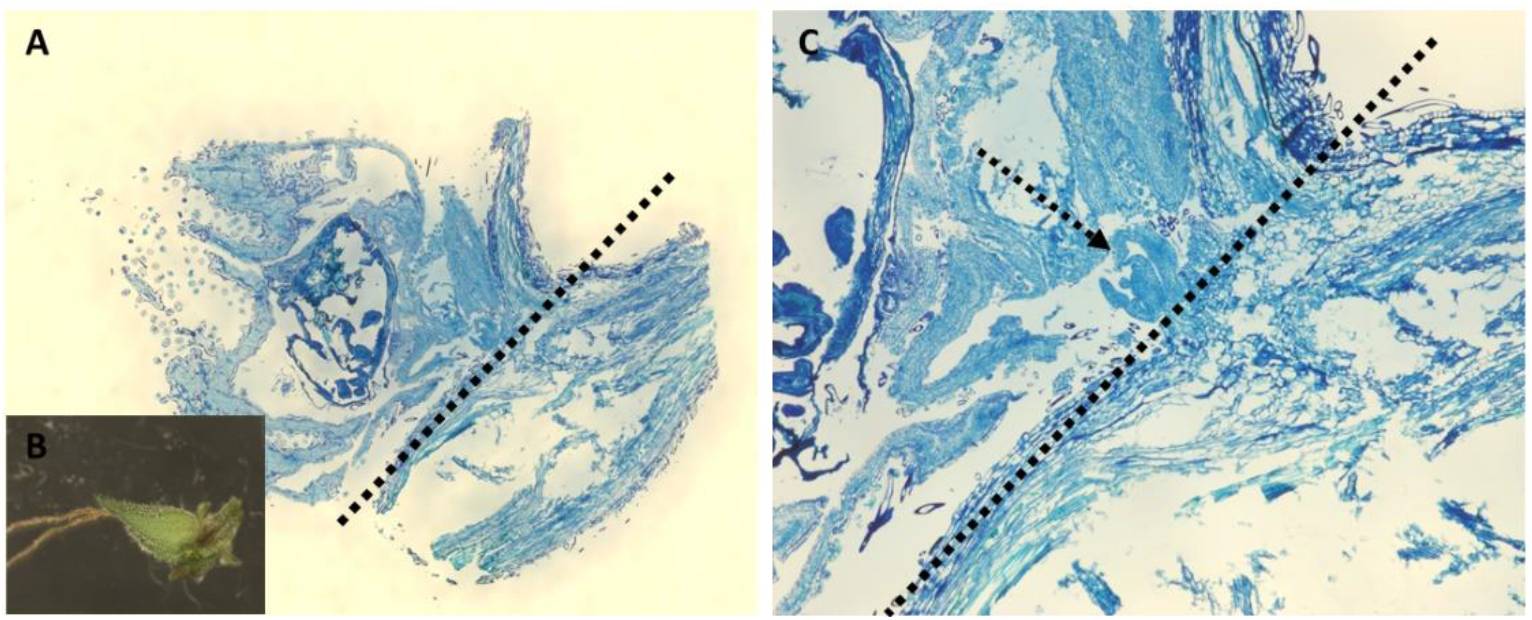
Cross-section of a single freshly dissected floret. A) Single freshly dissected floret. Dashed line shows the typical incision site when dissecting single florets. B) The same freshly dissection floret prior to embedding and histological sampling. C) Enlarged view the base of the floret shows a small meristem (dashed arrow) located at the base of the floret next to the typical excision site (dashed line).

**Fig. 8.**
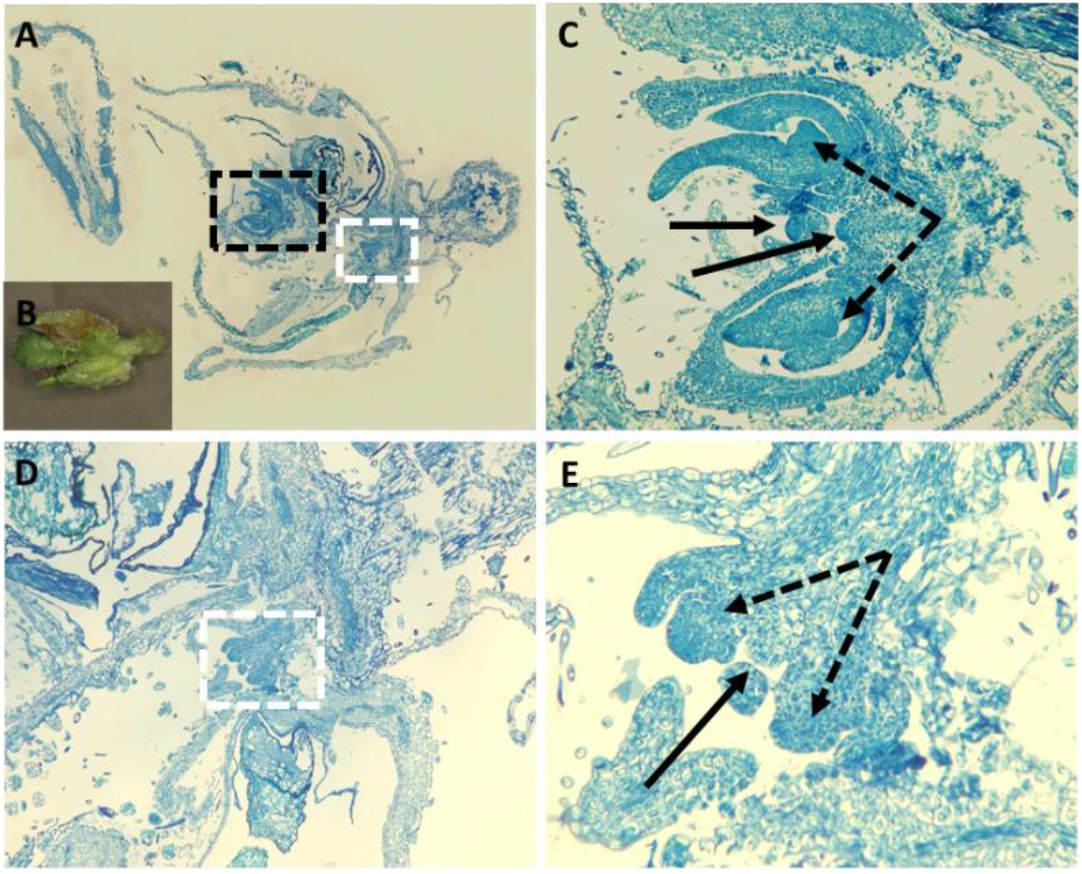
Pair of florets undergoing floral reversion 7 days post culture. A) Floral and vegetative meristem (dashed square) at the base of a pair of female *C. sativa* florets. B) The same 7-day-old floret prior to embedding and histological sampling. C) Enlarged view of the two floral meristems (dashed arrows) flanking two vegetative meristems (solid arrows) located at the base of excised florets from *C. sativa*. D) A second less developed cluster of immature vegetative and floral meristems highlighted by the white dashed lines, enlarged from A). E) Immature vegetative meristem (black solid arrow) flanked by two floral meristems (dashed arrows).

## Discussion

Floral reversion has been widely used for micropropagation of species recalcitrant to tissue culture, as well as in conservation efforts directed at threatened species (Appleton et al. 2012; Reshi et al. 2014; Kumar et al. 2015). This technique has achieved improved multiplication rates over conventional methods in important food crops such as banana, coconut and date palms, (Punyarani et al. 2013; Zayed et al. 2016; Shareefa et al. 2019). Floral reversion has not, however, been well explored in *C. sativa*, with only one published report of greenhouse derived floral tissue cultured on MS media supplemented with thidiazuron (TDZ; Piunno et al. 2019). In this work, Piunno *et al*. (2019) found that the reversion response was sporadic and only observed in 2/3 of cultivars tested. Additionally, though the cellular mechanism by which vegetative explants were produced was not explored, the authors hypothesized that reversion was occurring from existing meristems rather than de novo regeneration (Piunno et al. 2019). The present study aimed to expand upon these findings by using *in vitro* floral tissues and elucidating the cellular development leading to shoot production. We hypothesized that floral reversion using inflorescences from *in vitro* plants can provide an alternative approach to vegetative micropropagation of *C. sativa*.

In the study by Piunno *et al*. (2019), reversion was induced from clusters of florets varying in size from three to five florets. In the current study, we used single and pairs of florets rather than large clusters to increase the potential number of explants per inflorescence and to facilitate the subsequent histological study of the regenerating tissues. We found that pairs were 2.5-3 times more likely to revert than single florets in addition to being less labour-intensive to dissect than single florets. Our results indicate that floral reversion approaches can lead to significantly improved multiplication rates of 14.7 and 18.2 (from paired and single florets respectively) as compared to 2.2 plants produced from each nodal explant (Page et al. 2020). Lower multiplication figures reported by Page *et al*. (2020) using nodal tissues are likely impacted, in part, by the shorter timeframe of 35-43 days per cycle compared to the ~12 week timeframe for the proposed floral reversion cycle (Fig. 3) and future studies will help shorten the floral reversion timeframe. A notable reduction in labour can be achieved by using pairs of florets rather than single florets; which offer an only slightly higher multiplication index in return for increased labour. Our findings highlight the potential benefits of using floral reversion in a large-scale tissue culture setting, and we propose that floral reversion from floret clusters and investigation into automation of the explant preparation are pragmatic next steps in the optimization of this protocol for large-scale use.

Floral reversion has largely been shown to occur via organogenesis (Kavas et al. 2008; Appleton et al. 2012; Phulwaria and Shekhawat 2013; Punyarani et al. 2013; Poluboyarova et al. 2014; Kumar et al. 2015; Asker 2016); however, in some cases the mode of floral reversion remains unclear (Zayed et al. 2016; Shareefa et al. 2019). Floral reversion via indirect organogenesis has been demonstrated as a viable alternative to tradition culture methods in a range of species including *Cenchrus ciliaris* L. (buffel-grass)*, Arnebia hispidissima* (Lehm.) DC. (Arabian primrose), *Triticum durum* Desf. and *Triticum aestivum* L. (Kavas et al. 2008; Phulwaria and Shekhawat 2013; Kumar et al. 2015). In these species, floral reversion via indirect organogenesis required the inclusion of auxins and cytokinins for callus induction and subsequent regeneration of vegetative tissues. Inclusion of TDZ, which has both auxin and cytokinin-like activity (Guo et al. 2011), promoted callus induction in *C. sativa* inflorescences, but yielded low frequency of floral regeneration (Piunno et al. 2019). As a result, the current study focused on the proliferation of shoots using the cytokinins mT and BAP, rather than the induction of callus

Floral reversion has also been reported to occur via direct organogenesis through the formation of adventitious shoots and meristemoids from the base of the floral organs (Punyarani et al. 2013; Poluboyarova et al. 2014; Asker 2016). Histological sampling by Punyarani *et al*. (2013) revealed that shoot primordia with apical meristems, leaf primordia and procambium strands developing directly from the base of the floral explant in *Musa ssp*. Histological studies in *Allium altissimum* L. have also shown proliferation of shoots from the base of inflorescences with development of meristem centers occurring at the junction between the filament and tepal (Poluboyarova et al. 2014). Additional studies have suggested direct regeneration based on growth of shoots from the base of floral organs *in vitro*, but lack histological data validating these claims (Appleton et al. 2012; Asker 2016) and it is possible that reversion could be occurring from existing meristems in these cases.

In contrast our histological data provides evidence for the presence of an existing meristem region responsible for reversion at the base of freshly dissected florets. Histological sampling was carried out on freshly dissected florets as well as at the first signs of swelling and growth from single and pairs of florets at 7 days post culture. Sampling revealed the presence of a meristems at the base of the freshly dissected florets (Fig. 7) and histology of 7-day-old tissues reveals a distinct vegetative meristem flanked by floral meristems subtending the base of the ovary and the bracts (Fig. 8). We propose that this basal meristematic region is the genesis point for the observed shoot growth during floral reversion. Complete floral reversion, characterized by normal vegetative growth of the explant, appears to occur following a transitional period from the flowering stage to the ensuing vegetative growth stage during the first two weeks post-excision from the flowering *in vitro* mother plant. During this period, the newly excised florets would often produce one to two new florets along the elongating stem, prior to reverting to complete vegetative growth. This transitional growth can also be observed in whole plants grown *ex vitro* when a flowering plant is transferred to long days to “re-veg”, or when cuttings are taken from flowering plants. Histological samples from freshly dissected (day 0) florets reveal the existence of a meristematic region with apparent juvenile floral meristems; however no vegetative meristems could be distinguished (Fig. 7C).

We hypothesize that in this transition period existing proto-floral meristems continue development and reach maturity followed by the subsequent development of a vegetative meristem from the meristematic region at the base of the florets. The transition period ends with the change to vegetative growth from the meristem, typically within 2 weeks, thereby completing floral reversion. Alternatively, these meristems may develop adventitiously after excision from the flowering mother plant as a result in the disruption of the levels of endogenous plant growth regulators therefore triggering direct organogenesis. However, floral reversion occurred in the control treatments in the absence of plant growth regulators in this study, making *de novo* regeneration an unlikely candidate mechanism for floral reversion in *C. sativa*. The smaller percentage of responding explants observed in single florets may be attributed to damages sustained from dissection by the basal region housing the vegetative meristem responsible for shoot growth. In contrast, floret pairs which are held together by extra basal tissue following dissection, offer increased protection of this meristem region which consequentially led to higher percentage of reverting explants. As such we suggest that the lower response rates observed in single florets are due to the higher likelihood of a removal, or damage sustained by the meristems during the dissection process.

Meta-topolin has been reported to yield high percent responses and to promote multiple shoot formation in nodal multiple shoot cultures. Lata et al. (2016) found that mT supplementation between 1-4 μM resulted in 100% response rate and ~13 shoots per nodal explant. In the present study 1 uM mT achieved a maximum 81% reversion (pairs), despite producing fewer than 2 shoots per floral explant. These differences could be attributed to numerous factors, most importantly the difference in tissue source (*in vitro* florets vs. greenhouse-derived nodes). Responses to mT in *C. sativa* tissue culture have not, however, been universally successful. A recently published study by Wróbel et al. (2020) reported response rates of < 4% and shoot proliferation rates of <2 shoots/explant in response to mT treatment of nodal segments and shoot tips. Explants grown at 10 μM mT produced morphological abnormalities and showed signs of hyperhydricity, despite showing an increase in mass and canopy area over the 0 μM control in both singles and pairs. Morphological abnormalities attributable to hyperhydricity have been reported in *C. sativa* tissue cultures and have been suggested to be a result of nutrient imbalances and PGRs side-effects (Chaohua et al. 2016; Page et al. 2020). Despite low number of shoot proliferated from the florets, the potential number of explants that can be produced from a flowering *in vitro* explant is much higher than in a vegetative explant due to the floret-dense structure of the inflorescences (Spitzer-Rimon et al. 2019). In the present study, we used 1 μM mT treatments to calculate a floral multiplication index between 14.7 and 18.2, representing the estimated number of plantlets can be produced from a single *in vitro* flowering plant. These calculations highlight this method as a competitive option when compared with current nodal propagation methods. Improvements to the shoot proliferation stages through extensive screening of PGRs could make this method a vastly superior choice for many micropropagation needs when compared with contemporary nodal culture methods.

Piunno *et al*. (2019) reported low reversion rates despite the inclusion of TDZ in the medium, which has successfully promoted shoot proliferation in other *Cannabis* tissues as well as in many other species (Lata et al. 2009a, 2010; Monthony et al. 2020a). In the current study, we chose the synthetic cytokinin BAP as an alternative to TDZ. BAP is widely incorporated in floral reversion media (Eapen and George 1997; Phulwaria and Shekhawat 2013; Punyarani et al. 2013) and has been shown to promote shoot proliferation in non-floral tissues of other plant species (Jafari et al. 2011). In contrast with mT, the inclusion of BAP did not increase the canopy size, explant fresh weight, multiplication rate or the percentage of floret explants which reverted. While BAP alone may not enhance reversion, many studies report the use of BAP in combination with a second synthetic cytokinin and low levels of IAA (Eapen and George 1997; Gubišová et al. 2013; Reshi et al. 2014), and the use of a multi-PGR media for floral reversion in *C. sativa* is an important avenue for future exploration as this new *Cannabis* micropropagation technique is further developed.

In this study we elucidate the developmental origin of floral reversion in *C. sativa* and show that pairs of florets are more likely to undergo reversion than single dissected florets. In addition, we show that floral reversion improved our previously achieved multiplication rate using nodal cultures and we suggest that with further optimization of shoot proliferation and reduction of labour, floral reversion can exceed the potential for multiplication achieved using apical and axillary nodal cultures. We present the first histological evidence that reversion induced in *C. sativa* florets comes from existing meristems located in the floral tissues. We propose that the greater likelihood of reversion in pairs of florets is due to the vegetative meristems remaining intact during the dissection of pairs of florets. Our findings highlight the feasibility of developing floral reversion protocols and provide a roadmap for other groups wishing to assess floral reversion as viable alternatives to traditional vegetative micropropagation (Fig. 3). The exploration of the effects of cultivar, PGR and floret number presented herein will aid future researchers in the development of robust floral reversion protocols in the species.

## Acknowledgments

The authors gratefully acknowledge our industry partner, Hexo Corp. for the use of their plant material and Scott Golem for his help with greenhouse acclimatization of *in vitro* cultures. We would also like to thank Susan Lapos and the team at the Ontario Veterinary College’s Animal Health Laboratory for their histological expertise and support. The financial support of the Natural Sciences and Engineering Research Council of Canada (Grant No. RGPIN-2016-06252) is also gratefully acknowledged. Hexo Corp. (https://www.hexocorp.com) and the Natural Sciences and Engineering Research Council of Canada (https://www.nserc-crsng.gc.ca/index_eng.asp) were not involved in study design, data collection and analysis, the decision to publish or the preparation of the manuscript.

